# Extracellular Electron Transfer Enables Cellular Control of Cu(I)-catalyzed Alkyne-Azide Cycloaddition

**DOI:** 10.1101/2021.09.28.462180

**Authors:** Gina Partipilo, Austin J. Graham, Brian Belardi, Benjamin K. Keitz

**Affiliations:** McKetta Department of Chemical Engineering, University of Texas at Austin, Austin, TX, USA; Center for Dynamics and Control of Materials, University of Texas at Austin, Austin, TX, USA

## Abstract

Extracellular electron transfer (EET) is an anaerobic respiration process that couples carbon oxidation to the reduction of metal species. In the presence of a suitable metal catalyst, EET allows for cellular metabolism to control a variety of synthetic transformations. Here, we report the use of EET from the model electroactive bacterium *Shewanella oneidensis* for metabolic and genetic control over Cu(I)-catalyzed Alkyne-Azide Cycloaddition (CuAAC). CuAAC conversion under anaerobic and aerobic conditions was dependent on live, actively respiring *S. oneidensis* cells. In addition, reaction progress and kinetics could be further manipulated by tailoring the central carbon metabolism of *S. oneidensis*. Similarly, CuAAC activity was dependent on specific EET pathways and could be manipulated using inducible genetic circuits controlling the expression of EET-relevant proteins including MtrC, MtrA, and CymA. EET-driven CuAAC also exhibited modularity and robustness in ligand tolerance and substrate scope. Furthermore, the living nature of this system could be exploited to perform multiple reaction cycles without requiring regeneration, something inaccessible to traditional chemical reductants. Finally, *S. oneidensis* enabled bioorthogonal CuAAC membrane labelling on live mammalian cells without affecting cell viability, suggesting that *S. oneidensis* can act as a dynamically tunable biocatalyst in complex environments. In summary, our results demonstrate how EET can expand the reaction scope available to living systems by enabling cellular control of CuAAC.

## Introduction

Biological catalysis provides several advantages over traditional chemical catalysis including milder operating conditions, self-regeneration, and the ability to optimize activity via genetic manipulation^1-4^. Whole cell biocatalysts can leverage additional dynamic control over such reactions by coupling activity to cellular growth and metabolism^5^. However, reactions catalyzed by whole cells are typically limited to known metabolic transformations^6-7^. While efforts to augment the substrate scope of enzymatic reactions via directed evolution have been highly successful, there is still an ongoing need to expand the synthetic capabilities of live cells^7-8^.

Recently, we and others have connected cellular metabolism to exogenous synthetic reactions via hydrogen generation, the secretion of reactive cellular metabolites, and extracellular electron transfer (EET)^10-14^. Among these approaches EET is particularly advantageous because it provides a tunable protein bridge between central carbon metabolism and extracellular redox reactions, including those controlled via metal catalysts. Specifically, EET in the model electroactive bacterium *Shewanella oneidensis* (wild-type MR-1) is regulated through a set of well-defined heme-containing cytochromes in the Mtr-pathway (metal-reducing pathway)^15-17^. This pathway allows *S. oneidensis* to use oxidized metal ions, including organometallic catalysts, as terminal electron acceptors under anaerobic conditions^18-20^. Indeed, bacterial reduction of metals including iron(III) and copper(II) by *E. coli* and *S. oneidensis* have previously been used to perform atom-transfer radical polymerization (ATRP)^21-23^, and transcriptional regulation of specific EET proteins in *S. oneidensis* has enabled dynamic control over metal reduction and resulting catalysis^24-25^. Given these results we hypothesized that additional synthetic reactions involving the Cu(II/I) redox couple could be metabolically controlled using EET from *S. oneidensis*.

Cu(I)-catalyzed Alkyne-Azide Cycloaddition (CuAAC) is an example of bioorthogonal Click chemistry that exhibits fast kinetics and high specificity in complex environments^26-27^. The reaction involves a [2+3] cycloaddition between a terminal azide and terminal alkyne to create a product with a 1,4-disubstituted 1,2,3-triazole^28^. CuAAC has been used for almost two decades in applications including drug design, delivery, and synthesis^29-32^; polymer synthesis^33-35^; tissue engineering^35-37^; and bioorthogonal labelling^38-42^. As a result, expanding whole-cell catalysis to include this ubiquitous chemical transformation could yield significant control over triazole formation in unique environments. Because CuAAC is highly dependent on the oxidation state of Cu(II/I), we predicted it could be controlled via EET. Here, we co-opt EET from *S. oneidensis* to enable biological control over CuAAC in both anaerobic and aerobic environments. In contrast to previous examples of microbially assisted CuAAC^43-44^, we show the cycloadditions controlled by *S. oneidensis* are modulated through specific metabolic pathways and EET proteins, which can be manipulated through gene expression. Consistent with synthetic CuAAC, our microbial system was effective for a variety of different substrates and Cu(II/I) ligands. Unlike traditional chemical reductants, *S. oneidensis* cells enable multiple reaction cycles and dynamic CuAAC kinetics that are intrinsically linked to cellular metabolism. Finally, we demonstrate that EET-controlled CuAAC is effective in complex environments, such as a eukaryotic co-culture, without negatively impacting cell viability. Overall, our results highlight EET’s ability to enable non-enzymatic catalysis and place synthetic chemical reactions under living control.

## Results

### Extracellular electron transfer from live *S. oneidensis* catalyzes alkyne-azide cycloaddition via Cu(II) reduction

To monitor CuAAC reaction progress, we utilized a fluorogenic azide, CalFluor 488, which undergoes a Click-activated quenched-to-fluorescent shift^45^ (Figure 1a). This substrate allowed reaction progress to be monitored in real time with standard well plates. A typical EET-controlled CuAAC reaction consisted of CalFluor 488 and propargylated-PEG (Alkyne-PEG_4_-acid) with Cu(II) bromide and a Tris(benzyltriazolylmethyl)amine (THPTA) ligand in *Shewanella* Basal Medium (SBM). We initially compared anaerobic reaction conversion between wild-type *S. oneidensis* (inoculating OD_600_ = 0.1) and a chemical reducing agent, sodium ascorbate (NaAsc, 200 µM). After five hours, reactions containing NaAsc or *S. oneidensis* showed comparable conversion by fluorescence turn-on (Figure 1b). The presence of the desired triazole product was also confirmed with LC-MS (Figure S2). As expected, reactions lacking NaAsc, MR-1, or catalyst did not show any detectable conversion. While Cu(II/I) exhibits anti-microbial activity above certain concentrations, complexation to various ligands has been shown to mitigate the cytotoxic effects in *E. coli*^46^. Nevertheless, to account for the possibility of cell toxicity, we measured cell viability of *S. oneidensis* under typical reaction conditions containing 50 µM of CuBr_2_ and 300 µM of THPTA. Colony counting confirmed that CuAAC reagents did not cause cell death. Additionally, when employing a lower inoculating density (OD_600_ = 0.01) in equivalent reaction conditions, cells grew to anaerobic saturation (Figure S3). Finally, both mechanically lysed and heat-killed cells failed to achieve appreciable conversion, suggesting that actively respiring whole cells are required for the reaction to proceed. (Figure S4).

**Figure 1.**
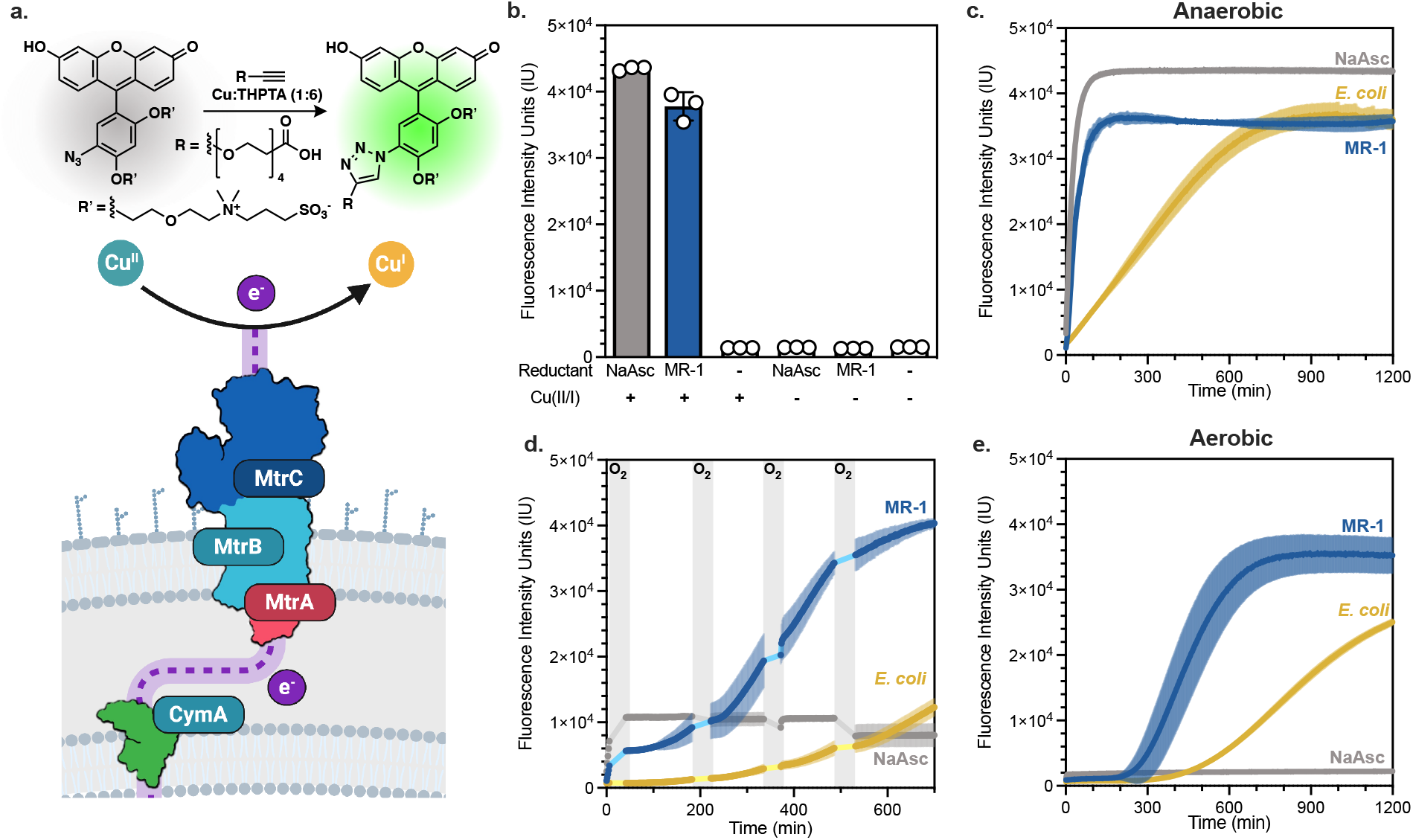
*S. oneidensis* enables both anaerobic and aerobic CuAAC. **a**. The Mtr-pathway shuttles electrons to the extracellular space, where they can reduce soluble Cu(II) *in situ* to Cu(I). The Cu(I) participates in CuAAC, yielding turn-on fluorescence upon cycloaddition (ex. 488 nm; em. 521 nm). **b**. CuAAC between 0.6 μM CalFluor 488 and 100 μM alkyne-PEG_4_-acid and 50 μM Cu:THPTA (1:6) in SBM. Anaerobic controls in presence or absence of Cu:THPTA and sodium ascorbate (NaAsc, 200 µM) or *S. oneidensis (*MR-1, OD_600_ = 0.1) measured after 5 hours. **c**. Anerobic kinetic curves monitoring the creation of the cycloaddition product in the presence of a sodium ascorbate (NaAsc, 200 μM), *S. oneidensis* (MR-1, OD_600_ = 0.1), or *E. coli* MG1655 (OD_600_ = 0.1). **d**. Kinetic curves monitoring anaerobic reactions exposed to oxygen (gray bars) before resealing and monitoring. **e**. Aerobic kinetic curves monitoring the creation of the cycloaddition product in the presence of sodium ascorbate (NaAsc, 200 μM), S. oneidensis (MR-1, OD_600_ = 0.1), or *E. coli* MG1655 (OD_600_ = 0.1). Data represent mean ± SD of n=3 biological replicates.

Next, we monitored CuAAC kinetics in real-time using fluorescence turn-on. Reactions containing wild-type *S. oneidensis* achieved comparable conversion to CuAAC using a chemical reductant over nearly identical time scales (Figure 1c). By comparison, wild-type *Escherichia coli* MG1655, which lacks specific EET proteins^47^, drove conversion more slowly than NaAsc or *S. oneidensis* (Figure 1c), consistent with previous evidence that non-specific Cu reduction is possible, but lacks comparable kinetics to EET^24-25^. Together, these results confirm that *S. oneidensis* MR-1 remains viable under our reaction conditions and can drive the CuAAC reaction via Cu(II) reduction.

### *S. oneidensis* catalyzes aerobic CuAAC without requiring dedicated oxygen removal

CuAAC often requires air-free conditions since Cu(I) is rapidly oxidized to Cu(II) in the presence of oxygen^38^. Because *S. oneidensis* is a facultative anaerobe and preferentially respires on oxygen, we hypothesized that our system could tolerate oxygen exposure and still drive CuAAC. To examine this possibility, anaerobically prepared reactions with either NaAsc, *S. oneidensis*, or *E. coli* were periodically unsealed, manually aerated, and left shaking under ambient conditions. Plates were then resealed, and the reaction was allowed to proceed. The aeration was repeated four times (Figure 1d). As expected, the reducing power of NaAsc was depleted after the first aeration event. Similarly, *E. coli*, which exhibited significant background activity under anaerobic conditions, was hindered by the repeated Cu(I) oxidation and was unable to achieve substantial conversion. In contrast, *S. oneidensis* drove the reaction to completion despite repeated oxygen exposure, presumably consuming dissolved oxygen through aerobic respiration before resuming EET^25^. Next, we challenged *S. oneidensis* to perform microbial CuAAC with an aerobic pre-growth and benchtop set up. Under these conditions, *S. oneidensis* achieved comparable conversion to anaerobic reactions (Figure 1e). Standard concentrations of NaAsc failed to achieve any significant conversion aerobically and reactions with *E. coli* were again significantly arrested. In fact, greater than 500 μM of sodium ascorbate was required to achieve any notable aerobic conversion and greater than 1000 μM, a 20-times excess relative to Cu(II), was required to achieve comparable conversion to *S. oneidensis* (Figure S6). In contrast to chemical reductants, the lag time between cell inoculation and reaction turn-on could be modulated by changing the initial density of *S. oneidensis* (Figure S6). Together, these results indicate that *S. oneidensis* is unique in its ability to sustain CuAAC conversion under benchtop conditions by upregulating EET and Cu(II) reduction in response to oxygen depletion.

### Observed rate of copper reduction can be modeled with simple reactions

To compare the kinetics of microbial CuAAC, we developed a simplified reaction model that accounts for oxygen consumption, copper reduction/oxidation, and cycloaddition. This model facilitated parameter estimation from time course kinetics by monitoring the concentration of the triazole product (Equation S1, Figure 2a). We calculated the observed parameter fitting for rate constants for our standard *S. oneidensis* CuAAC system from the following simplified reactions (Equation S2-S5).

**Figure 2.**
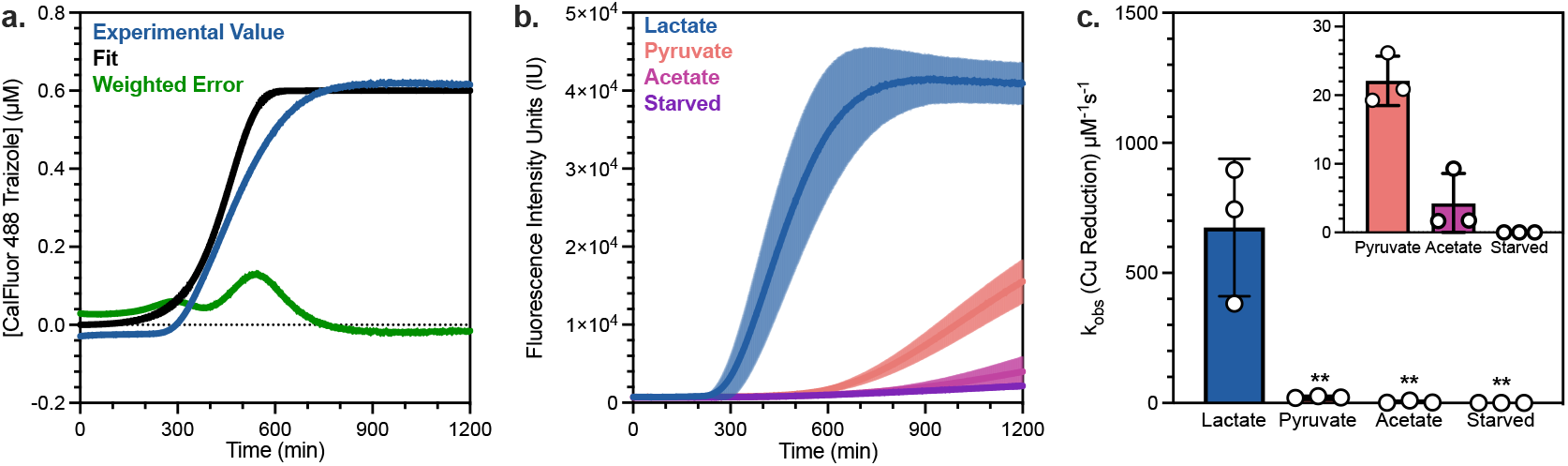
Kinetic modeling enables differentiation of microbial CuAAC reactions through manipulating central carbon metabolism. **a**. Modeling data (black) compared to the average of n=18 aerobic *S. oneidensis* MR-1 (OD_600_ = 0.1) (blue) with weighted error (green). **b**. Aerobic kinetic curves with varying carbon source identities (20 mM). **c**. Quantification of the observed rate of copper reduction (k_obs_ Cu Reduction, μM^-1^ s^-1^) for varying carbon sources (20 mM) performed by rate fitting kinetic curves. Data show mean ± SD of n=3 biological replicates. **P<0.01

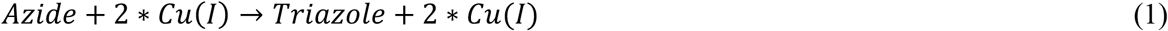

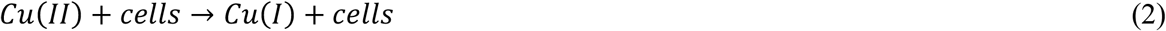

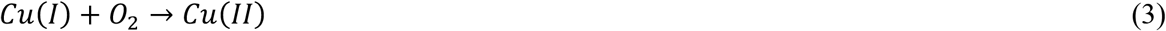

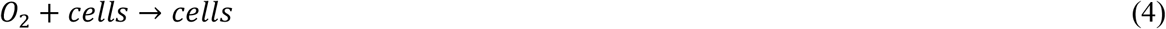

After fixing all other observed reaction rate constants, we performed a parameter estimation on the triazole formation curves to determine the rate of copper reduction (k_obs_ Cu Reduction μM^-1^s^-1^). In cases where cell growth was relevant to the parameter estimation, as occurs with a lower starting inoculum, the observed rate of cell growth was also fit using cell growth curves which were experimentally monitored through absorption at 600 nm. The agreement between the fit and the experimental data is outlined in Figure 2a. This simple model provides a convenient handle for quantifying and comparing kinetic rate differences between various reaction conditions and strains.

### Carbon metabolism controls the rate of *S. oneidensis* catalyzed CuAAC

The requirement for viable cells and the relationship between inoculating density and reaction lag time in our aerobic reactions suggested that cellular respiration influences conversion and that manipulating carbon metabolism may exert control over the reaction. Anaerobic carbon metabolism in *S. oneidensis* is well-defined; it cannot utilize acetate as a carbon source under these conditions but generates four electron equivalents per molecule of lactate and two electron equivalents per molecule of pyruvate^48-49^. To test the effect of carbon source on conversion, microbial CuAAC reactions were performed in reactions that contained either lactate, pyruvate, or acetate as the carbon source (Figure 2b). As expected, the cycloaddition reaction proceeded at the greatest rate when cells were grown on lactate (k_obs_ = 674 ± 264 μM^-1^ s^-1^). With starved or acetate-fed cells, the reaction was significantly attenuated. Consistent with our hypothesis, pyruvate-fed cells proceeded with a greater lag time and slower rate (k_obs_ = 22 ± 3 μM^-1^ s^-1^) compared to lactate (Figure 2c). These results indicate that live cell carbon metabolism is critical for both aerobic respiration and copper reduction through EET and that the CuAAC reaction rate can be directly controlled by the carbon source provided to *S. oneidensis*.

### Specific Mtr-pathway proteins control the rate of *S. oneidensis* catalyzed CuAAC

In the absence of oxygen, *S. oneidensis* expresses the cytochromes in the CymA/Mtr-pathway (Figure 1a)^50^. Electrons are transported via the cytoplasmic membrane protein, CymA, which reduces periplasmic proteins that provide electrons to the Mtr-pathway. Electrons are then shuttled through MtrA and onto MtrC, where they are deposited onto a terminal electron acceptor^16,51^. Given the importance of the Mtr-pathway in regulating EET, we tested a series of EET-deficient strains for their ability to perform aerobic CuAAC. As expected, the observed reaction rate significantly decreased upon the removal of *mtrC* (Figure 3a-b). Additionally, removal of *mtrF*, which encodes an MtrC homologue, further decreased the observed reaction rate from 402 ± 66 μM^-1^ s^-1^ to 193 ± 33 μM^-1^ s^-1^. Finally, complete removal of the Mtr-pathway caused the reaction kinetics to closely resemble the background reduction observed in *E. coli*. Similar to wild-type *S. oneidensis*, the lag time of the knockouts was also dependent on inoculating density. Lag times for reactions involving *E. coli* did not show a similar dependence (Figure S7). Together, these results confirm that the rate of CuAAC is primarily controlled by EET via the Mtr-pathway.

**Figure 3.**
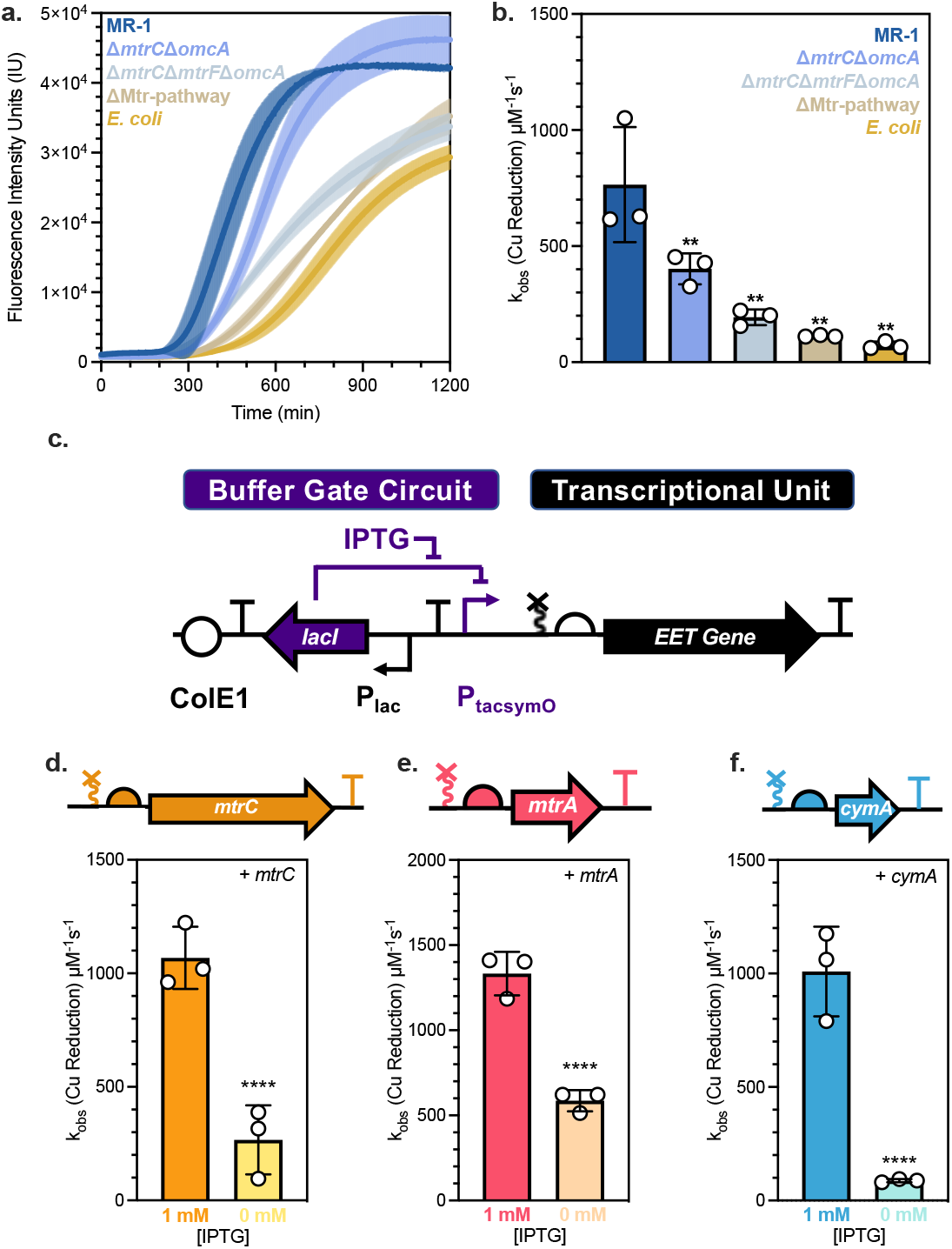
Control over CuAAC using genetic engineering of the CymA/Mtr-pathway. **a**. Aerobic kinetic curves performed with various *S. oneidensis* knockouts (OD_600_ = 0.1) and an *E. coli* MG1655 negative control. **b**. Quantification of the observed rate of copper reduction for varying *S. oneidensis* knockouts or an *E. coli* negative control. **c**. Diagram of generic Buffer gate circuit used to control expression of the gene of interest (*mtrC, mtrA*, or *cymA*) in *S. oneidensis* knockout strains^18^. Quantification of the observed rate of copper reduction for raw kinetic curves inoculated at an OD_600_ = 0.025 (5 × 10^7^ CFU/mL) with **d**. JG596 + *mtrC*, **e**. Δ*mtrA + mtrA*, and **f**. Δ*cymA + cymA* for fully induced and uninduced constructs. Data show mean ± SD of n=3 biological replicates. **P<0.01, ****P<0.0001

### Transcriptional regulation of the Mtr-pathway enables controllable CuAAC

Our results using EET-deficient knockouts suggested control of CuAAC rates and conversion could be achieved using genetic engineering. Thus, we aimed to control CuAAC activity via inducible transcription of EET genes. Specifically, we used a *S. oneidensis* Δ*mtrC*Δ*omcA*Δ*mtrF* strain harboring *mtrC* on a plasmid under control of the *P*_*tac*_ promoter^52^ (Figure 3c). The signaling molecule isopropyl ß-D-1-thiogalactopyranoside, IPTG, activates *mtrC* gene expression, which we predicted would regulate CuAAC rate and conversion. All genetic constructs tested were grown overnight in the absence of IPTG and induced upon inoculation into CuAAC reaction mixtures (Figure 3d). At the same initial inoculum and in the presence of 1 mM IPTG, CuAAC reaction kinetics using a complemented *mtrC* knockout in a Δ*mtrC*Δ*omcA*Δ*mtrF* strain closely resembled those of wild-type *S. oneidensis*. However, conversion kinetics resembled those of the parent-knockout strain in the absence of IPTG (Figure S10 and S11). Reactions using empty vector controls in both wild-type *S. oneidensis* and Δ*mtrC*Δ*omcA*Δ*mtrF* closely agreed with their parent strains. (Figure S8).

We similarly regulated two other EET-relevant genes, *mtrA* and *cymA*, in their cognate knockouts and saw comparable differences in observed rate constants in response to IPTG (Figure 3e-f). For each of these inducible constructs, the observed rate of Cu-reduction was rescued by the complementation of the cognate gene. To further emphasize the degree of transcriptional control over CuAAC activity, we fit reaction conversion using inducible gene expression models for each strain (Figure S11). Overall, these results confirm that catalyst reduction is driven by EET and can be placed under transcriptional control.

### Substrate tolerance of microbially catalyzed CuAAC

Having demonstrated that CuAAC can be controlled through EET, we proposed that modulating the copper reduction potential through complexation to various stabilizing ligands could further tune reaction kinetics, similar to chemical catalysis (Figure 4a). Specifically, we found that BTTAA (2-(4-((bis((1-(tert-butyl)-1H-1,2,3-triazol-4-yl)methyl)amino)methyl)-1H-1,2,3-triazol-1-yl)acetic acid) exhibited higher activity, consistent with literature reports^53^, in the presence of *S. oneidensis* as well as in NaAsc controls (Figure S12). When the kinetics for each copper stabilizing ligand were modeled, we observed that the rate of copper reduction when utilizing BTTAA (1895 ± 339 μM^-1^ s^-1^) was significantly faster than that of THPTA (585 ± 188 μM^-1^ s^-1^) (Figure 4b), broadly reflecting the observed time course data. Tris(2-pyridylmethyl)amine (TPMA) is a ligand often used for atom-transfer radical polymerization (ATRP) and is reported as being ineffective for the CuAAC reaction^54^. Consistent with these results, TPMA did not show any significant conversion and yielded similar results as the ligand-free system under our conditions. These results indicate that an appropriate ligand is required for significant CuAAC activity and that *S. oneidensis* CuAAC conforms to known synthetic CuAAC-ligand trends.

**Figure 4.**
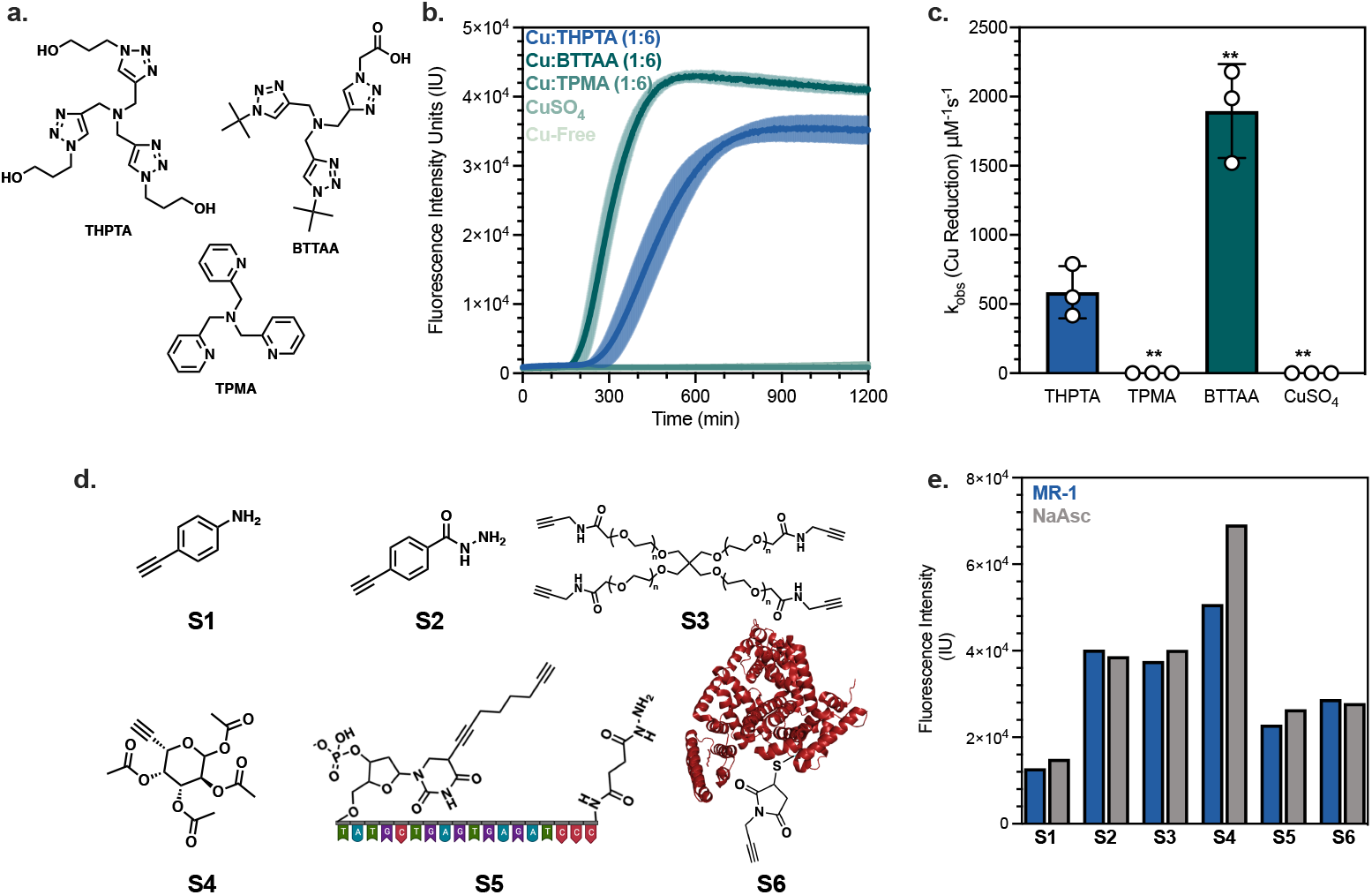
The effect of copper ligand on *S. oneidensis* catalyzed CuAAC and accessible substrate scope. **a**. Chemical structures for THPTA, BTTAA, and TPMA. **b**. Aerobic kinetic curves for CuAAC between 0.6 μM CalFluor 488 and 100 μM alkyne-PEG_4_-Acid in SBM with varying copper ligand identities in a 1:6 ratio (50μM:300μM). **c**. Quantification of the observed rate of copper reduction (k_obs_ Cu Reduction, μM^-1^ s^-1^) for varying copper ligand sources performed by rate fitting kinetic curves. Data show mean ± SD of n=3 independent experiments. **P<0.01 **d**. Various alkyne identities Top: 4-ethynylaniline (100 µM), 4-ethynylbenzohydrazide (100 µM), 4-arm-PEG (10k) (25 µM), Bottom: Click It™ fucose (100 µM), ssDNA oligio (20 bp) with terminal alkyne (100 µM), BSA-alkyne functionalized at C35 (83 µM, 1 mg/mL). **e**. Fluorescent conversion monitoring CalFluor 488 triazole formation with various alkyne substrates with *S. oneidensis* MR-1 and sodium ascorbate (NaAsc, 1 mM).

In contrast to most enzymatic reactions, EET acts on a soluble metal catalyst, which can then react with a broad range of potential substrates. We examined a series of alkyne-containing substrates including small molecules (4-ethynylaniline and 4-ethynylbenzohydrazide), a sugar (Click-IT™ Fucose Alkyne), a 4-arm long chain polymer (5k), a functionalized single-stranded DNA, and a protein (bovine serum albumin, BSA) for their ability to undergo CuAAC in the presence of *S. oneidensis*. Regardless of alkyne identity, *S. oneidensis* successfully performed the reaction aerobically with comparable conversion to positive controls using NaAsc (Figure 4e).

### Microbial CuAAC enables repeated cycling utilizing adherent *S. oneidensis* cells

A significant advantage of our system is the regeneration capability of bacterial cells and the potential to repeatedly perform CuAAC reactions (Figure 5a). To demonstrate this, we first conducted a CuAAC reaction during which *S. oneidensis* adhered to the bottom of a Nunc-coated 96-well plate (Figure 5b). After completion of the reaction, the product-containing supernatant was decanted and replaced with fresh starting material, leaving the adhered cells intact. Each cycle of the system yielded consistent conversion (Figure 5c, Figure S13) and cycles III and IV reflect remarkably similar reaction rates to the initial reaction. The increase in observed rate for cycle II likely reflects an increase in cell inoculum from a partial transition to aerobic respiration during oxygen exposure and corresponding growth. In subsequent cycles, the cells appear to perform the reaction at the same rate as the initial turnover. To further demonstrate the robustness of the system and highlight the capability to tune reaction kinetics *in situ*, the carbon source provided to *S. oneidensis* was changed between cycles, alternating between pyruvate and lactate. Cycles with *S. oneidensis* grown on pyruvate maintained microbial viability but did not display significant conversion over the course of the 11-hour cycle. When the reaction material was replaced with a solution containing lactate, the CuAAC conversion increased (Figure 5d, Figure S13). This OFF/ON cycling highlights how *S. oneidensis* can dynamically control CuAAC by sensing and reacting to changes in reaction conditions.

**Figure 5.**
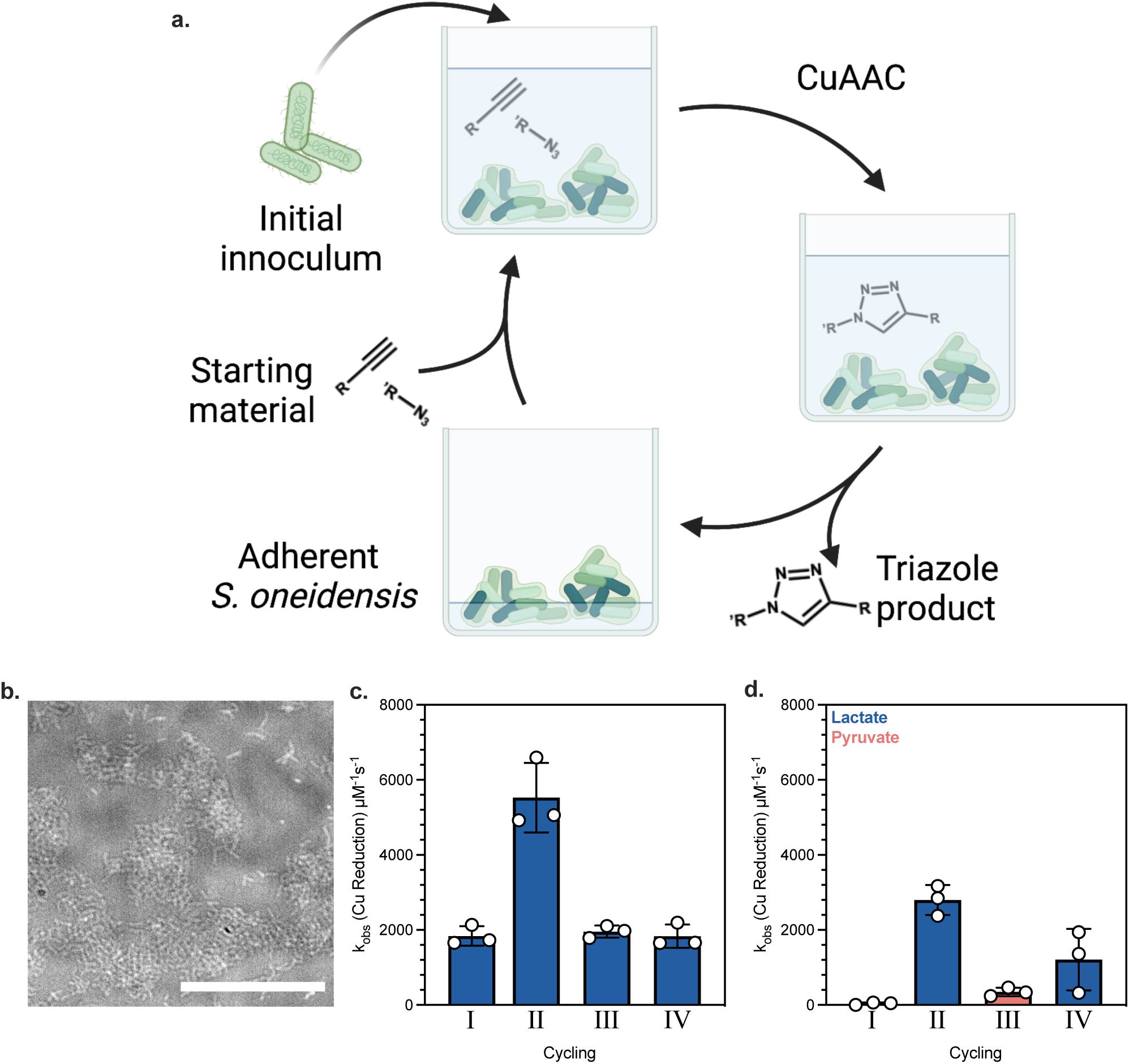
Adherent *S. oneidensis* allows for repeated cycling of CuAAC without regeneration. **a**. Cycling experimental set up. **b**. Representative bright field image of *S. oneidensis* cells adhered to Nunc-coated 96-well plate after removal of reaction supernatant. Scale bar represents 50 µm. **c**. Repeat CuAAC kinetics utilizing the same batch of *S. oneidensis* for each 11-hour cycle. **d**. Repeat CuAAC kinetics utilizing the same batch of *S. oneidensis* with different carbon sources for each 11-hour cycle. Data show mean ± SD of n=3 biological replicates.

### *S. oneidensis* enables CuAAC in Mammalian Co-Culture

CuAAC is notable for its orthogonality in complex biological settings and bacteria are well suited for long-term or responsive applications in these environments. However, for applications involving eukaryotic organisms, it is critical that *S. oneidensis* controls CuAAC without affecting eukaryotic viability. To assess this, we first performed CuAAC with our standard azide and alkyne partners in a 3T3 murine fibroblast and *S. oneidensis* co-culture. Under our conditions, *S. oneidensis* was as effective at performing CuAAC as 1 mM of chemical reductant NaAsc (Figure 6a). After 90 minutes, neither the CuAAC reactants nor the bacteria negatively impacted viability (Figure 6b). A microscopy time-series revealed that bacteria remained distributed and motile in co-culture (Supplementary Video 1). Together, these results indicate that *S. oneidensis* can exert dynamic control over CuAAC in mammalian co-culture without diminishing viability.

**Figure 6.**
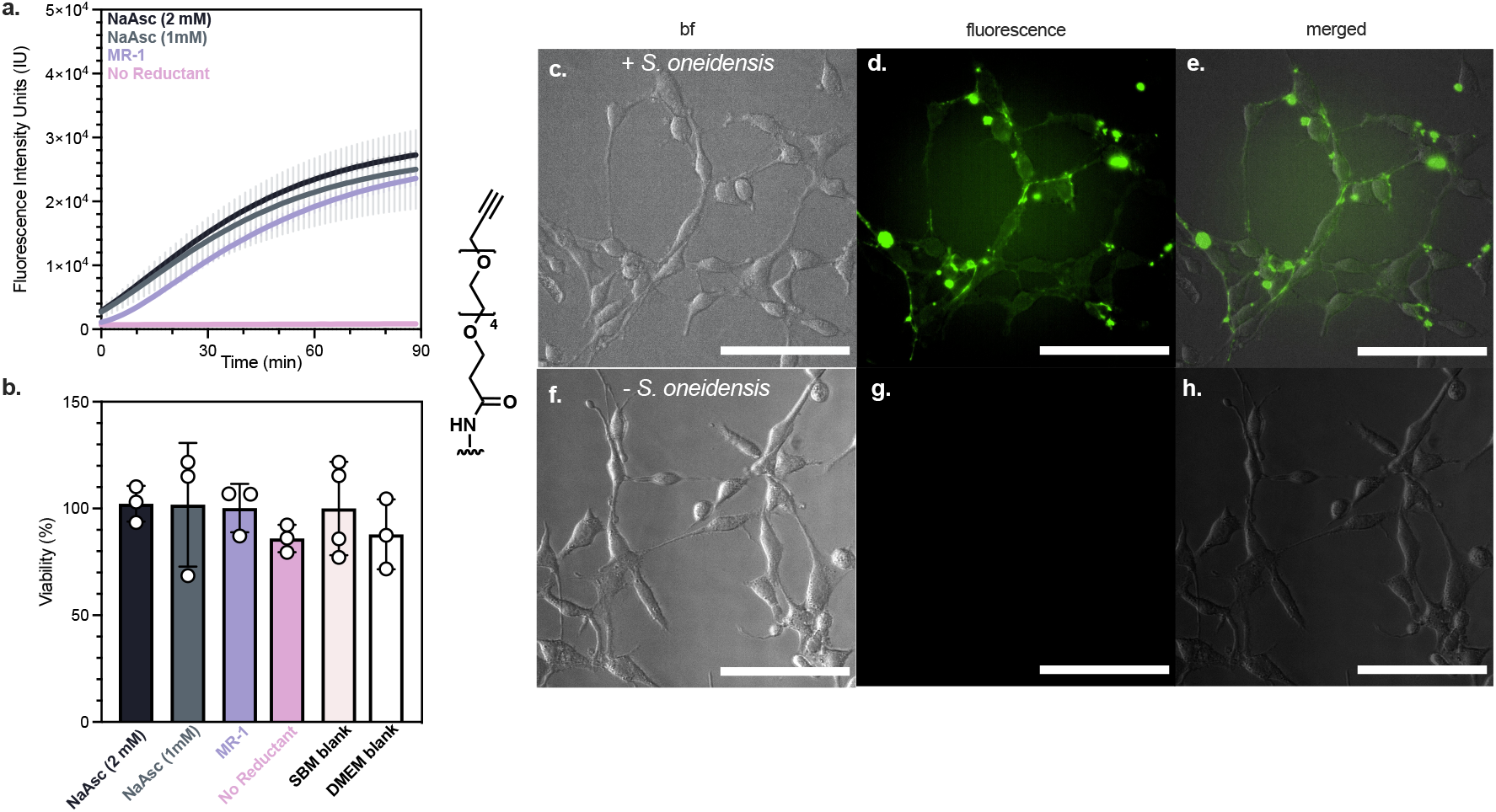
CuAAC performed in the presence eukaryotic cells. **a**. Kinetic curves monitoring the creation of the cycloaddition product in the presence of sodium ascorbate (NaAsc, 1 mM and 2 mM), *S. oneidensis* (MR-1, OD_600_ = 0.1), or cell-free controls. **b**. Cell viability post-CuAAC reaction in Figure 6a measured via MTT assay. All results are normalized by SBM blank. Data show mean ± SD of n=3 biological replicates. **c–h**. Surface functionalization with PEG-alkyne through NHS-ester displacement by free amines reacted for 3 h with CalFluor 488 in the presence of *S. oneidensis:* **c**. bright field image, **d**. fluorescent image, and **e**. merged image and in the absence of *S. oneidensis*: **f**. bright field image, **g**. fluorescent image, and **h**. merged image. Scale bars indicate 100 µm.

Next, *S. oneidensis* was used to label the membranes of fibroblast cells. Fibroblast cells were NHS-ester functionalized with alkyne-PEG_4_. Following esterification, cells were washed and CalFluor 488 probe, Cu(II), and THPTA were added along with *S. oneidensis* to the reaction vessel. After completion of the reaction, fibroblast cells were washed and imaged using epifluorescence microscopy. Increased fluorescence intensity along the cell membranes confirmed formation of the triazole product on mammalian cell surfaces (Figure 5c-e). In a bacteria-free control, no notable fluorescence could be detected, indicating that the reaction was dependent on the presence of *S. oneidensis* (Figure 5f-h). In a subsequent experiment to demonstrate the modularity of the system, fibroblast cells were functionalized with 6-azidohexanoic acid (a terminal azide), and the CuAAC reaction was successfully performed with a carboxyrhodamine 110 terminal alkyne probe (Figure S14). Together, these results suggest that *S. oneidensis* CuAAC is compatible with mammalian cells and can potentially be applied in traditional CuAAC settings including biorthogonal labelling^55-56^, -omics^38, 57-58^, and tissue engineering^59-60^.

## Discussion

Overall, we successfully developed a whole-cell microbial redox biocatalyst for small-molecule CuAAC Click reactions. Employing EET for non-enzymatic conversion significantly expands the substrate scope available to bacteria and facilitates genetic and metabolic control over this important chemical transformation. We first leveraged *S. oneidensis* as a dynamic actuator for controlling anaerobic and aerobic CuAAC reactions. Our results highlight how the central metabolism of *S. oneidensis* can be manipulated to control reaction lag time, kinetics, and conversion. Similarly, after repeated oxygen exposures, we showed that our system was less susceptible to oxygen challenges compared to traditional chemical reductants. Furthermore, transcriptionally regulating *mtrC* and other EET genes increased kinetic control and conversion. Thus, changes to central metabolism could be used in tandem with genetic engineering techniques to modulate the reaction kinetics over several orders of magnitude (Figure S7 and S9).

In negative controls involving the ΔMtr-pathway knockout or *E. coli*, we did not observe a complete arrest of CuAAC conversion. It is likely that Cu(II) reduction is tied to other electron transport or reduction pathways such as extracellular flavins^61^, glutathione^14,43,62^, and copper nanoparticle formation^63^. Despite the presence of some background reduction, our results indicate that conversion is primarily controlled by the Mtr-pathway. Nevertheless, a key focus of future work will be fine tuning dynamic range and increasing genetic control over the reaction while mitigating background reduction. Potential solutions are to limit reduction of extracellular copper solely to the Mtr-pathway and could include expression of the Mtr-pathway in non-native host organisms^64^, creation of flavin exporter knockouts^61^ or decreasing glutathione production^43^.

We demonstrated substrate robustness of *S. oneidensis* CuAAC by successfully reacting an alkyne-functionalized sugar, nucleic acid, protein, and several small molecules. In all cases conversion was comparable to treatment with NaAsc. Additionally, adhered *S. oneidensis* performed aerobic CuAAC over multiple cycles. In these experiments, the bacteria did not require regeneration or intervention even after several repeated oxygen exposures. In contrast to traditional chemical or biological reductants (e.g. NaAsc or NADH), CuAAC activity could be dynamically tuned by interchanging simple carbon sources. Changes to the central carbon metabolism of *S. oneidensis*, such as engineered glucose catabolism, could be further utilized to tune CuAAC kinetics^65^. Together, our results demonstrate that *S. oneidensis* is comparable to traditional CuAAC reductants but with added benefit of dynamic metabolic and genetic regulation.

Finally, in complex environments such as mammalian cell culture, *S. oneidensis* enabled CuAAC without impacting cell viability. While more investigations are needed to determine the full effect of *S. oneidensis* on mammalian cells, our successful co-culture experiments lay the foundation for applying our system toward traditional CuAAC applications such as drug delivery^66^, cell encapsulation^37^, tissue scaffolding^55^, cell labelling^45-46^, non-canonical amino acid incorporation^67^, and more^37^. Excitingly, several of these applications could benefit from the genetic, metabolic, and temporal control available to our system. In summary, our results demonstrate how EET combines the advantages of biological control and synthetic catalysis to expand the chemical reaction space available to microbes.

## Materials

CalFluor 488 (Click Chemistry Tools), Alkyne-PEG_4_-Acid (Click Chemistry Tools), Copper(II) bromide (CuBr_2_, Sigma-Aldrich, 99%), tris(2-pyridylmethyl)amine (TPMA, Sigma-Aldrich, 98%), Alkyne Tris(benzyltriazolylmethyl)amine (THPTA, Sigma-Aldrich, 95%), 2-(4-((bis((1-(tert-butyl)-1H-1,2,3-triazol-4-yl)methyl)amino)methyl)-1H-1,2,3-triazol-1-yl)acetic acid (BTTAA, Click Chemistry Tools > 95%), Click-IT™ Fucose Alkyne (Invitrogen), 6-Azidohexanoic Acid Ester (Click Chemistry tools, > 95%), Alkyne-PEG_4_-NHS Ester (Click Chemistry Tools > 95%), Bovine serum albumin alkyne (Click Chemistry Tools), sodium DL-lactate (NaC_3_H_5_O_3_, TCI, 60% in water), sodium fumarate (Na_2_C_4_H_2_O_4_,VWR, 98%), HEPES buffer solution (C_8_H_18_N_2_O_4_S, VWR, 1 M in water, pH = 7.3), potassium phosphate dibasic (K_2_HPO_4_, Sigma-Aldrich), isopropyl ß-D-1-thiogalactopyranoside (IPTG, Teknova), potassium phosphate monobasic (KH_2_PO_4_, Sigma-Aldrich), sodium chloride (NaCl, VWR), dimethyl sulfoxide (cell culture grade, Sigma-Aldrich), kanamycin sulfate (C18H38N4O15S, Growcells) ammonium sulfate ((NH_4_)_2_SO_4_, Fisher Scientific), magnesium(II) sulfate heptahydrate (MgSO_4_·7H_2_O, VWR), trace mineral supplement (ATCC), casamino acids (VWR), DMEM high glucose pyruvate (Life Technologies), Fetal Bovine Serum (Life Technologies), Trypsin-EDTA solution (Sigma-Aldrich), CyQUANT MTT Cell Viability Assay (Life Technologies), penicillin-streptomycin solution (Sigma-Aldrich), silicone oil (Alfa Aesar) were used as received. All media components were autoclaved or sterilized using 0.2 μm PES filters.

## Methods

### Analysis and Measurement

Fluorescence emission was collected on a BMG LABTECH CLARIOstar plate reader with a 491(± 14) nm and an emission collection at 538 (± 38) nm). After the addition of all plate components, the 96-well plate was sealed with a sterile and optically transparent sealing film (PCR-SP-S, AxySeal Scientific) and covered with a polystyrene plate lid (Eppendorf) lined with silicone grease. The plate reader was held at 30 °C and collected emissions every 90 seconds for 10 to 24 hours.

### Microscopy

All microscopy was performed using a Nikon Ti2 Eclipse inverted epifluorescence microscope. Fluorescence was measured using a GFP excitation/emission filter cube on the Nikon Ti2.

### Bacteria Strains and Culture

Bacterial strains and plasmids are listed in Table S1. Cultures were prepared from bacterial stocks stored in 20% glycerol at −80 °C streaked onto LB agar plates (for wild-type and knockout strains) and grown overnight at 30°C for *Shewanella* and 37 °C for *E. coli*. Overnight cultures were grown by picking single colonies and inoculating into *Shewanella* basal medium (SBM) (Table S2) supplemented with 0.05% w/v trace mineral supplement, 0.05% w/v casamino acids, and 20 mM sodium lactate (2.85 μL of 60% w/w sodium lactate per 1 mL culture) as the electron donor. Aerobic cultures were grown in 15 mL culture tubes at 30 °C and 250 rpm shaking. Anaerobic cultures were grown using the same procedure, but in argon sparged growth medium supplemented with 40 mM sodium fumarate as the electron acceptor in a Coy Anaerobic Glovebox containing a humidified atmosphere at 3% hydrogen content and the balance nitrogen. Plasmid harboring strains were grown with the addition of 25 μg/mL of kanamycin diluted from a 1000× stock in water. Cultures were washed 3× after overnight growth using SBM supplemented with 0.05% casamino acids (degassed for anaerobic cultures)^23^. OD_600_ was measured using a NanoDrop 2000C spectrophotometer and normalized to an OD_600_ of 0.75 before dilution into reaction mixture. All CuAAC reactions used 26.7 μL of OD_600_ = 0.75 concentrated cell culture into 173.3 μL of reaction mixture to give a final OD_600_=0.1, ca. (1.8 ± 0.5) × 10^8^ CFU·mL^-1^, unless otherwise noted.

### Standard Microbial CuAAC

All reactions were performed in *Shewanella* basal media (SBM) supplemented with 0.05% w/v casamino acids with lactate (20 mM) as a carbon source, and fumarate (20 mM) as the primary electron acceptor. Stock solutions of 1 M sodium fumarate and 60 w/v% lactate solutions were stored at 4°C until use. Aliquots of 1.2 mM of CalFluor 488^45^ were created in DMSO and stored frozen at -80 °C until use. Aliquots of 4 mM Alkyne-PEG_4_-Acid were created in DMSO and stored at -20 °C until use. An 8 mM copper bromide stock in DMF was created and stored at 4 °C and mixed with an equal volume amount of 48 mM freshly made stock of THPTA in sterile water. In alternative copper ligand studies, a 48 mM solution of the ligand in water or methanol was mixed with copper bromide. In order, the following was added to yield a 200 μL reaction in either degassed or ambient SBM supplemented with 0.05% casamino acids with the final concentrations: lactate (20 mM) (or alternative carbon source), fumarate (20 mM), Cu:THPTA 1:6 (50 μM: 300 μM)^68^, Alkyne-PEG_4_-Acid (100 μM), CalFluor 488 (0.6 μM), and finally *S. oneidensis* (OD_600_ of 0.1) or freshly dissolved NaAsc in water (200 μM). The reaction was then placed into the plate reader for analysis and allowed to react for between 10 and 24 hours.

### Microbial CuAAC Controls

Heat-killed controls were obtained by incubating bacterial cultures (post-wash) at an OD_600_ of 0.75 at a temperature of 80 °C for 15 minutes^24^. Upon completion, the cells solution was vortexed to ensure complete mixing, and diluted (26.7 μL into 173.3 μL) into the reaction mixture. Mechanically lysed cells were obtained via sonication with a Branson Model 250 sonicator with a Model 102C Converter. Cells suspensions at an OD_600_ of 0.75 were placed on ice and sonicated at 30% strength for 2.5 minutes with cycles of 10 seconds and 5 seconds between cycles. This process was repeated 3×. Upon completion, the transparent solution was vortexed to ensure complete mixing, and diluted (26.7 μL into 173.32 μL) into the reaction mixture.

### Oxygen Exposure CuAAC

A standard anaerobic microbial CuAAC reaction was begun utilizing either NaAsc (200 μM), *S. oneidensis* (OD_600_ = 0.1) or *E. coli* (OD_600_ = 0.1). The reaction was sealed and allowed to progress while monitoring the fluorescent output for 6 minutes before removing the lid and aerating by bubbling 10 mL of ambient air over a 30 second period into each 200 μL reaction. The reactions were then shaken for 20 minutes, without a lid, and re-sealed to begin collecting fluorescence for two hours. This was repeated four times.

### Cycling experiments CuAAC

A standard aerobic microbial CuAAC reaction was begun in a Nunclon Delta-Treated 96-well plate (Thermo Scientific 167008). After sealing and allowing to react for 11 h, the supernatant was removed carefully, as to not disturb the layer of cells on the bottom of the well. Starting materials in fresh SBM (200 µL) was gently added back into the well, the plate was sealed and again allowed to react for 11 h. This was repeated three times for a total of four reactions.

### Mammalian Cell Culture

3T3 fibroblast cells (American Type Culture Collection, gifted from the Rosales Lab at UT Austin) were cultured in T75 flasks between 9-13 passages in Dulbecco’s modified Eagle’s medium (DMEM) supplemented with 10% fetal bovine serum (FBS) and 1% penicillin-streptomycin. Cells were cultured in a humidified incubator at 37 °C with 5% CO_2_. At ∼80% confluence, the cells were washed with PBS buffer, cleaved with trypsin solution (0.25% trypsin containing 0.02% ethylenediamine tetraacetic acid in PBS), centrifuged, and seeded at a new confluency of 20%.

### MTT assay

3T3 cells were used for assays after the 9th passage and before the 13th passage. The cells were plated at 4 × 10^4^ cells/well in a Nunclon Delta-Treated 96-well plate (Thermo Scientific 167008). The cells were allowed to adhere overnight, the supernatant was aspirated, and the cells were washed with 1X PBS. Then, 173.3 µL of the appropriate reaction mixture or control was added to the wells and the reaction was initiated through the addition of 26.7 µL of *S. oneidensis* (OD_600_=0.75) grown from an anaerobic overnight or appropriate media blank. Control wells contained only 200 µL of SBM + 0.05% w/v casamino acids. A DMEM control was also run to ensure there was no detriment to viability resulting from SBM. The plate was then sealed as described previously and after a 1.5 h incubation, the plates were removed from the plate reader and imaged for 30 minutes. At this time the supernatant was aspirated and then, 100 µL of DMEM and 10 µL of CyQUANT MTT Cell Viability Assay in 1X PBS (Thermo Scientific V13154) was added. After a 4-hour incubation with the MTT the supernatant was aspirated, and the wells were resuspended in 100 µL of dimethyl sulfoxide (DMSO). Absorption measurements were collected at 490 nm using the plate reader.

### Cell Surface Functionalization

3T3 cells used for assays after the 9th passage and before the 13th passage. The cells were plated at 2 × 10^4^ cells/well in a Nunclon Delta-Treated 96-well plate (Thermo Scientific 167008). The cells were allowed to adhere overnight, the supernatant was aspirated, and the cells were washed with 1X PBS (pH 7.4) to remove any proteins from the culture media. A 100 µM solution of the desired NHS-Ester in PBS (pH 7.4) was created fresh from a 4 mM stock of the NHS-Ester in DMSO. The solution was added (200 µL) to each well and allowed to react at room temperature for 30 minutes. The supernatant was then removed, and each well washed with PBS (pH 7.4). To each reaction vessel a solution of: corresponding probe (0.6 µM), lactate (20 mM), fumarate (20 mM), Cu:THPTA 1:6 (50 μM: 300 μM), finally *S. oneidensis* (OD_600_ of 0.1) in SBM supplemented with 0.05% w/v casamino acids was added. The reactions were allowed to incubate for 3 h at 30 °C and washed with PBS (pH 7.4) before being taken to the Nikon Ti2 Eclipse inverted epifluorescence microscope. Images were taken using a 1 s exposure time using a GFP channel.

### Observed Rate of Cu-Reduction

Utilizing COmplex PAthway SImulator (COPASI) each of the proposed reactions (Reactions 1-4) was input into the biological model^69^. All fluorescence measurements were converted to CalFluor 488 triazole concentration using the calibration curve outlined in Figure S1. Fitting Reaction 1 first with kinetic data collected from an anaerobic CuAAC using sodium ascorbate, and under the assumption that the triazole formation is the limiting step^45^, the k_1-obs_ was obtained through parameter estimation and was fixed. Next, k_2-obs_ was obtained by fitting Reaction 2 with kinetic data collected from an anaerobic CuAAC using *S. oneidensis* (Figure S5) and was fixed. Finally, the observed rate constants for Reaction 3 and 4 were fit simultaneously using parameter estimation and kinetic data from aerobic CuAAC using *S. oneidensis*. The corresponding observed rate constants for each reaction were fixed and are outlined in Table S4. Each k_obs_ (Cu-Reduction) was obtained by un-fixing k_2-obs_ and performing a parameter estimation with the fixed rate constants. As this crude model aims to only quantify the observed rate kinetics, it is assumed that only the rate of Cu-Reduction is changed when changes are made to EET (knockouts, various carbon sources, complementation, etc.).

### Statistical Analysis

Unless otherwise noted, data are reported as mean ± SD of *n* = 3 biological replicates. Significance was calculated in GraphPad Prism 9.0 using a two-tailed unpaired student *t* test or a one-way ANOVA.

## Author Contributions

G.P., A.J.G., and B.K.K. conceived the project. G.P. performed experiments. G.P. and A.J.G. performed microscopy. B.B. and B.K.K. supervised research. G.P., A.J.G., and B.K.K. analyzed the results. All authors wrote the manuscript.

## Acknowledgements

This work was financially supported by the Welch Foundation (Grant F-1929, B.K.K.), the National Institutes of Health under award number R35GM133640 (B.K.K.), an NSF CAREER award (1944334, B.K.K.), and the Air Force Office of Scientific Research under award number FA9550-20-1-0088 (B.K.K.). G.P. and A.J.G were supported through National Science Foundation Graduate Research Fellowships (Program Award No. DGE-1610403). *S. oneidensis* knockouts (JG596, JG1194, JG749) were a generous gift from Prof. Jeffry Gralnick (U. Minnesota). 4-arm-PEG-Alkyne was a generous gift from Prof. Adrianne M. Rosales (UT Austin) and Thomas M. FitzSimons. Small molecule alkynes (4-ethynylaniline and 4-ethynylbenzohydrazide) were a generous gift from Prof. Eric V. Anslyn (UT Austin) and Dr. Cecil J. Howard. *E. coli* MG1655 was a generous gift for Prof. Lydia Contreras (UT Austin). Plasmid-harboring *S. oneidensis* strains were created by Dr. Christopher M. Dundas and used with his gracious permission. Mass spectrometry were kindly performed and interpreted by Dr. Ian Riddleton and Dr. Kristin Blake. Elements of Figure 1, Figure 4, and Figure 5 were created with BioRender.com.

