## Supplementary Information for "Extracellular Electron Transfer Enables Cellular Control of Cu(I)-catalyzed Alkyne-Azide Cycloaddition"

**Table S1.** Strains, plasmids, and DNA used in this study

| Strain or Plasmid | Description/Genotype | Reference/Source |
|---|---|---|
| ***S. oneidensis* Strains** | | |
| MR-1 | MR-1 (ATCC700550), wild-type strain | American-Type Culture Collection |
| JG749 | Lacks outer membrane cytochromes, MtrC and OmcA; ΔmtrCΔomcA | [1] |
| JG596 | Lacks out membrane cytochromes MtrC, OmcA, and MtrF; ΔmtrCΔomcAΔmtrF | [1] |
| JG1194 | Lacks numerous proteins responsible for EET, including outer membrane cytochromes, βbarrel cytochromes, and periplasmic electron carriers; ΔMtr | [2] |
| MR-1+pCD8 | Wild-type with empty Buffer Gate | [3] |
| Δ*cymA*+pCD26r1 | Δ*cymA* with *cymA* Buffer Gate | [3] |
| Δ*mtrA*+pCD25r0 | Δ*mtrA* with *mtrA* Buffer Gate | [3] |
| JG596+pCD24r1 | JG596 with *mtrC* Buffer Gate | [3] |
| JG596+pCDd1 | JG596 with *mtrC* NOT Gate | [3] |
| JG596+pCD8 | JG596 with empty Buffer Gate | [3] |
| ***E. coli* Strains** | | |
| MG1655 | Wild-type strain | Lydia Contreras, U. of Texas at Austin |
| **Eukaryotic Cells** | | |
| 3T3 | Fibroblast cells, American Type Culture Collection | Adrianne Rosales, U. of Texas at Austin |
| **ssDNA** | **sequence** | |
| ssDNA-Alkyne | /5ILink12/CCC TAG AGT GAG TCG TAT G/35OctdU/ | IDT |

1. Coursolle, D; Gralnick, J.A. Modularity of the Mtr respiratory pathway of Shewanella oneidensis strain MR-1. *Molecular Microbiology*. **2010**, *77*, 995–1008.

2. Coursolle, D; Gralnick, J.A. Reconstruction of extracellular respiratory pathways for iron(III) reduction in *Shewanella oneidensis* strain MR-1. *Front. Microbiol.* **2012,** *3* (2), 1–11
3. C. M. Dundas, D.J.F Walker, B. K. Ketiz. Tuning Extracellular Electron Transfer by *Shewanella oneidensis* Using Transcriptional Logic Gates. *ACS Synth. Bio*. **2020**, *9*(9), 2301-2315.

**Table S2.** Ingredients in *Shewanella* Basal Medium. Growth media was supplemented with casamino acids and Wolfe's mineral solution. Reaction media was only supplemented with casamino acids.

| Ingredient | Quantity for 1 L of 1 X SBM |
|---|---|
| $K_2HPO_4$ | 225 mg |
| $KH_2PO_4$ | 225 mg |
| NaCl | 460 mg |
| $(NH_4)_2SO_4$ | 225 mg |
| $MgSO_4 \cdot 7H_2O$ | 117 mg |
| HEPES | 100 mL of 1 M HEPES |
| Casamino acids | 5 mL of 10% casamino acids |
| Wolfe's mineral solution | 5 mL of Wolfe's Mineral Solution |
| $ddH_2O$ | Up to 1 L, adjust to pH = 7.2 |

**Conversion:**

$$Emission_{528} = 50{,}252 * [Triazole\ (\mu M)] + 2{,}297 \qquad \text{(S1)}$$

**Observed Rate Constant Equations:**

$$\frac{d([Azide])}{dt} = -k_{CuAAC}[Azide][Cu(I)]^2 \qquad \text{(S2)}$$

$$\frac{d([Triazole])}{dt} = k_{CuAAC}[Azide][Cu(I)]^2 \qquad \text{(S3)}$$

$$\frac{d([Cu(I)])}{dt} = k_{Cu-reduction}[Cu(II)][Cells] - k_{Cu-oxidation}[Cu(I)][O_2] \qquad \text{(S4)}$$

$$\frac{d([Cu(II)])}{dt} = -k_{Cu-reduction}[Cu(II)][Cells] + k_{Cu-oxidation}[Cu(I)][O_2] \qquad \text{(S5)}$$

$$\frac{d([O_2])}{dt} = -k_{Cu-oxidation}[Cu(I)][O_2] - k_{respiration}[O_2][cells] \qquad \text{(S6)}$$

**Table S4.** Observed Rate Constants ($k_{obs}$) as determined by COPASI

| Constant | Value | Units |
|---|---|---|
| $k_{1-obs}$ | $4.03 \times 10^{-6}$ | $\mu M^{-2}\, s^{-1}$ |
| $k_{2-obs}$ (initial value) | 1398.54 | $\mu M^{-1}\, s^{-1}$ |
| $k_{3-obs}$ | $3.53 \times 10^{-6}$ | $\mu M^{-1}\, s^{-1}$ |
| $k_{4-obs}$ | 672.59 | $\mu M^{-1}\, s^{-1}$ |

**Modeling of Inducible Constructs**

Standard CuAAC reaction was performed in the presence of various amounts of IPTG inducer molecule (1000 μM, 500 μM, 100 μM, 50 μM, 25 μM, 10 μM, 1 μM, and 0 μM) and inoculated with a bacterial cell stock of the plasmid-containing strains at a normalized $OD_{600}$ of 0.2 for dilution into reaction mixture. 26.7 μL of the concentrated cell culture was added into 173.3 μL of reaction mixture to give a final $OD_{600}$=0.025, ca. $(4.5 \pm 0.5) \times 10^7$ CFU·mL$^{-1}$. The fluorescence readings were converted to conversion utilizing the calibration curve in Figure S1. Utilizing GraphPad Prism 9.0 the reaction conversion was modeled as an activating Hill functional y = min + (max – min) $\frac{[I]^n}{{K_{1/2}}^n+[I]^n}$. The fluorescence readings were plotted versus corresponding IPTG values. Fitting parameters and "goodness of fit" can be found in Table S3.

**Table S3.** Hill Function constants

| Strain | Min (μM) | Max (μM) | Hill Slope (n) | $K_{1/2}$ (μM) | Goodness of fit |
|---|---|---|---|---|---|
| pCD24r1 | 0.44 | 0.82 | 1.464 | 33.57 | 0.8021 |
| pCD25r0 | 0.26 | 0.58 | 0.8501 | 169.7 | 0.4220 |
| pCD26r1 | 0.28 | 0.71 | 0.8234 | 28.92 | 0.8234 |

### **Supplementary Figures**

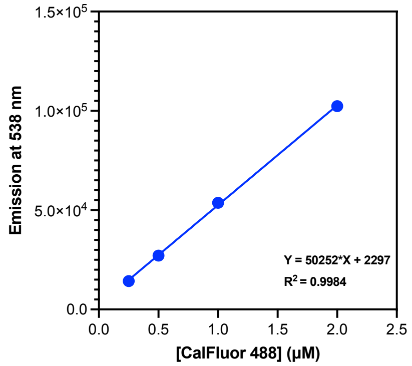


**Figure S1.** Calibration curve for CalFluor 488 triazole product after CuAAC with Alkyne-PEG$_4$-acid. Each point indicates the average n=3 ± standard deviation.

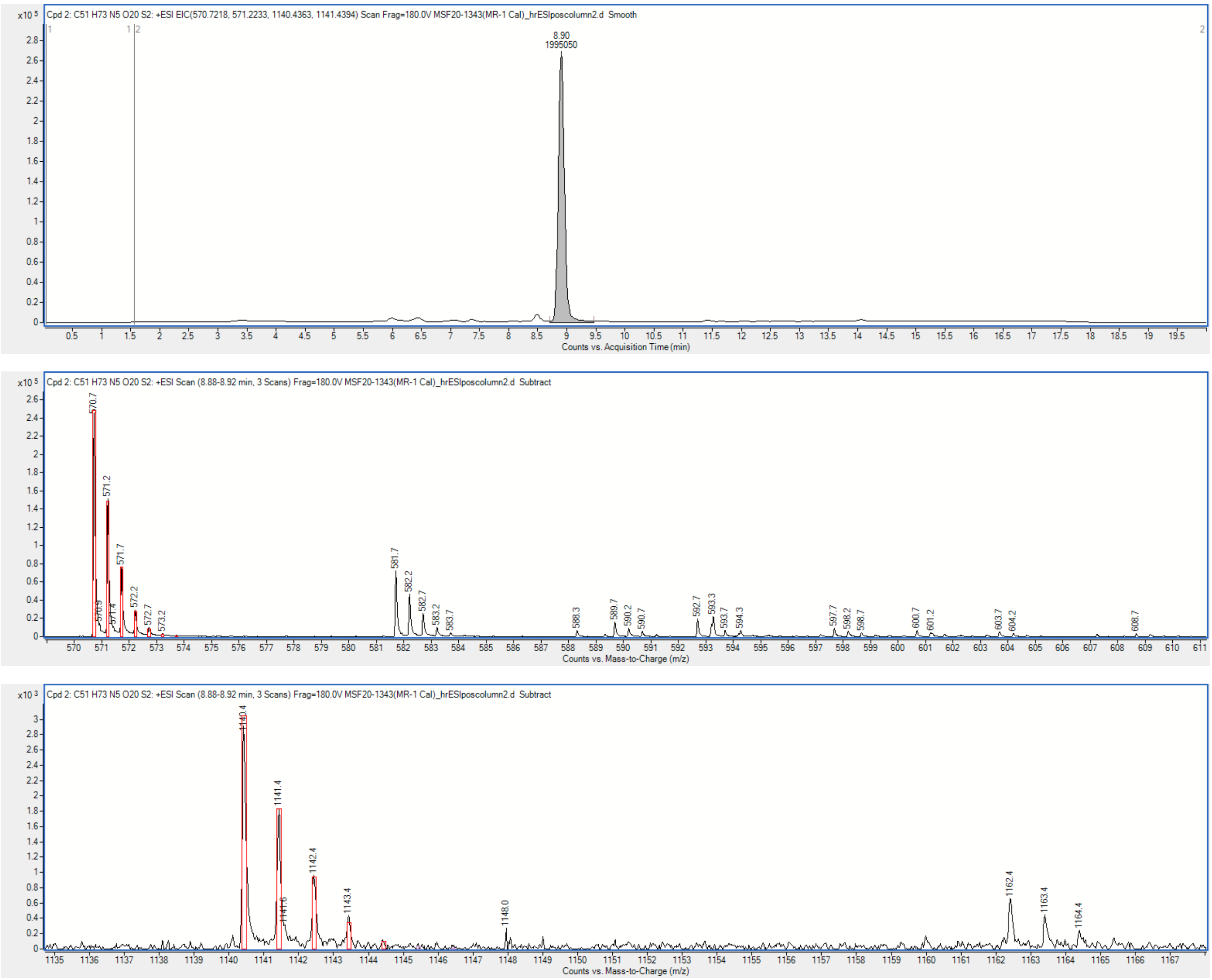


**Figure S2.** LC MS results for CalFluor 488 triazole product (exact mass 1139.34 g/mol) after a 20-hour CuAAC reaction with *S. oneidensis*. No starting material detected.

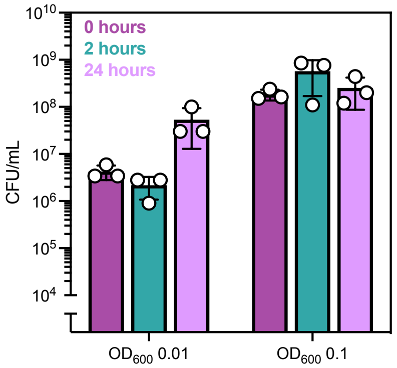


**Figure S3.** Colony counts for standard CuAAC reaction (inoculating density $OD_{600} = 0.1$) and decreased inoculum ($OD_{600} = 0.01$) to determine toxicity after 2 and 24 hours. Data show mean ± SD of n=3 replicates.

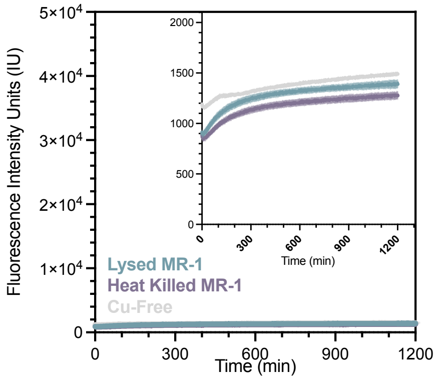


**Figure S4.** CuAAC between 0.6 μM CalFluor 488 and 100 μM alkyne-$PEG_4$-Acid in a SBM in the presence of mechanically lysed and heat killed *S. oneidensis* or in Cu-free conditions. Data show mean ± SD of n=3 replicates.

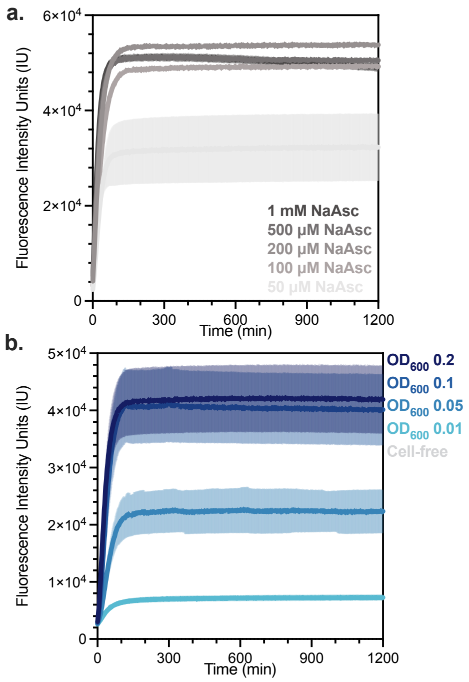


**Figure S5. a.** CuAAC between 0.6 µM CalFluor 488 and 100 µM alkyne-PEG4-Acid in SBM. Anaerobic reaction kinetics in the presence of various amounts of sodium ascorbate (NaAsc 1 mM, 500 µM, 200 µM, 100 µM, 50 µM) or **b.** different inoculating density of *S. oneidensis* ($OD_{600}$ 0.2, 0.1, 0.05, 0.01 or cell-free). Data show mean ± SD of n=3 replicates.

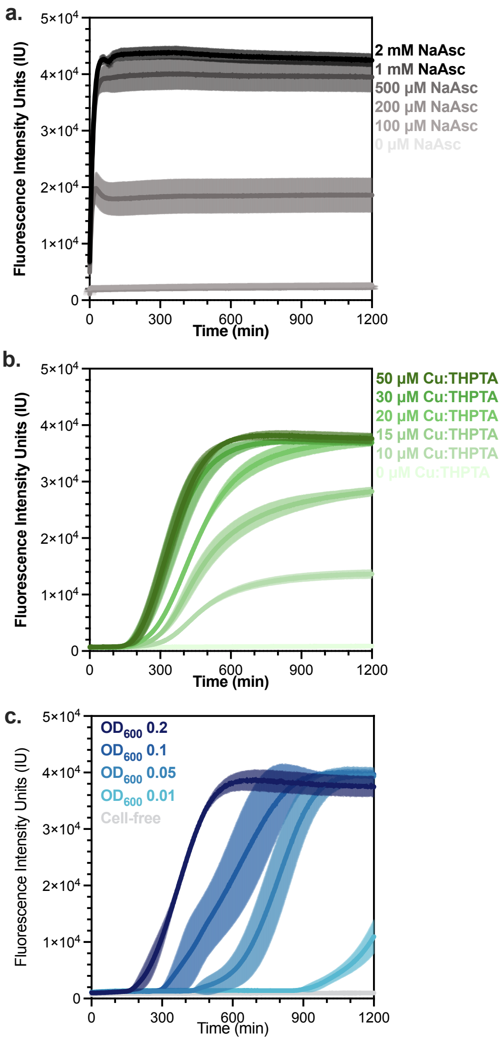


**Figure S6. a.** CuAAC between 0.6 µM CalFluor 488 and 100 µM alkyne-PEG$_4$-Acid in SBM. Aerobic reaction kinetics in the presence of various amounts of sodium ascorbate (NaAsc 2 mM, 1 mM, 500 µM, 200 µM, 100 µM, 0 µM), **b.** utilizing different amounts of copper catalyst ($CuBr_2$ 50 µM, 30 µM, 20 µM, 15 µM, 10 µM, 0 µM) complexed to a THPTA ligand in a ratio of 1:6 ($CuBr_2$:THPTA) or **c.** different inoculating density of *S. oneidensis* ($OD_{600}$ 0.2, 0.1, 0.05, 0.01 or cell-free). Data show mean ± SD of n=3 replicates.

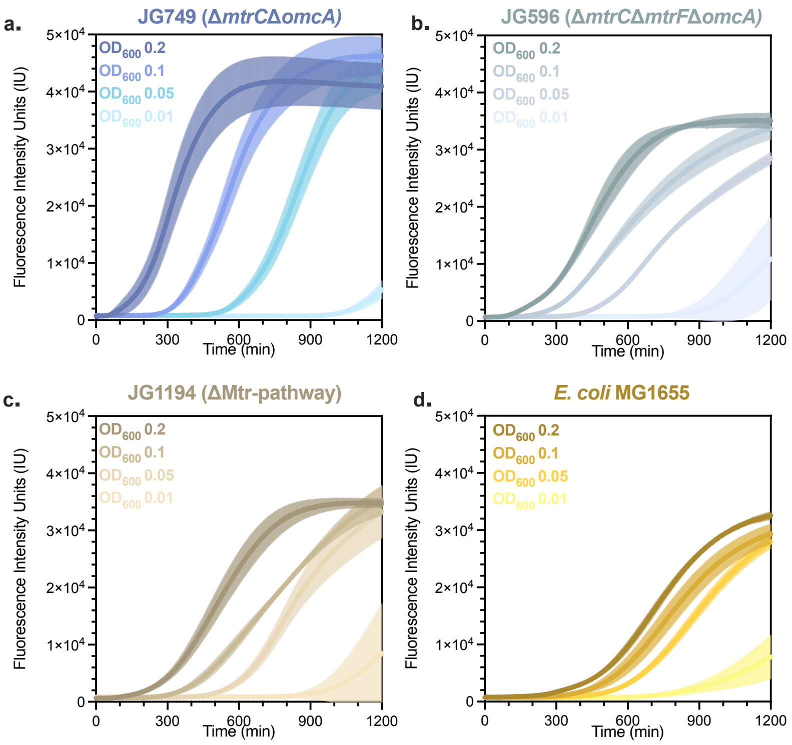


**Figure S7.** CuAAC between 0.6 μM CalFluor 488 and 100 μM alkyne-PEG$_4$-Acid in SBM. Aerobic reaction kinetics with various inoculating densities of **a.** *S. oneidensis* JG749 (Δ*mtrC*Δ*omcA*), **b.** JG596 (Δ*mtrC*Δ*omcA*Δ*mtrF*), **c.** JG1194 (ΔMtr-pathway) and **d.** *E. coli* MG1655. Data show mean ± SD of n=3 replicates.

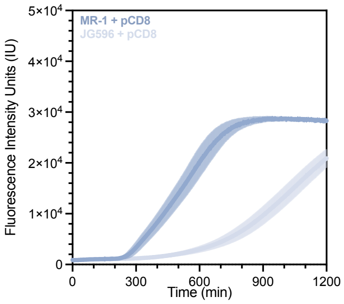


**Figure S8.** CuAAC between 0.6 μM CalFluor 488 and 100 μM alkyne-PEG$_4$-Acid in SBM. Aerobic reaction kinetics utilizing empty plasmid backbones (pCD8) in *S. oneidensis* MR-1 and JG596 (Δ*mtrC*Δ*omcA*Δ*mtrF*). Data show mean ± SD of n=3 replicates.

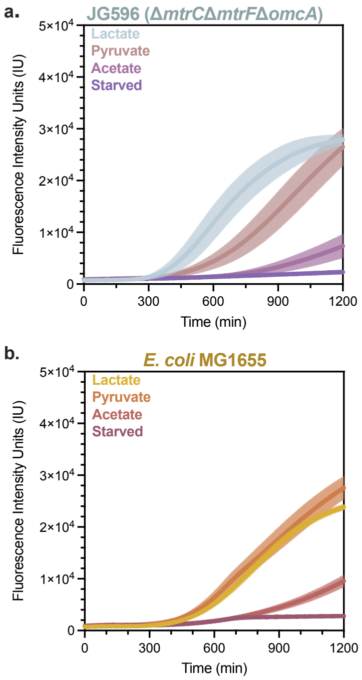


**Figure S9.** CuAAC between 0.6 μM CalFluor 488 and 100 μM alkyne-PEG$_4$-Acid in SBM. Aerobic reaction kinetics utilizing various carbon sources with **a.** *S. oneidensis* JG596 (Δ*mtrC*Δ*omcA*Δ*mtrF*) and **b.** *E. coli* MG1655. Data show mean ± SD of n=3 replicates.

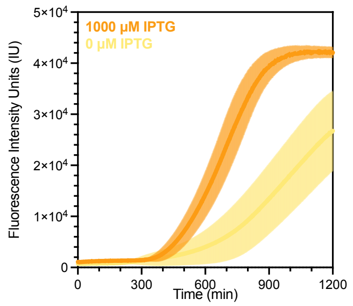


**Figure S10.** CuAAC between 0.6 μM CalFluor 488 and 100 μM alkyne-PEG4-Acid in SBM. Raw aerobic kinetic curves for JG596 + pCD24r1 with *mtrC* inducible plasmid for both fully induced and uninduced kinetics inoculated at an $OD_{600}$ = 0.1 (1.8 x $10^8$ CFU/mL). Data show mean ± SD of n=3 replicates.

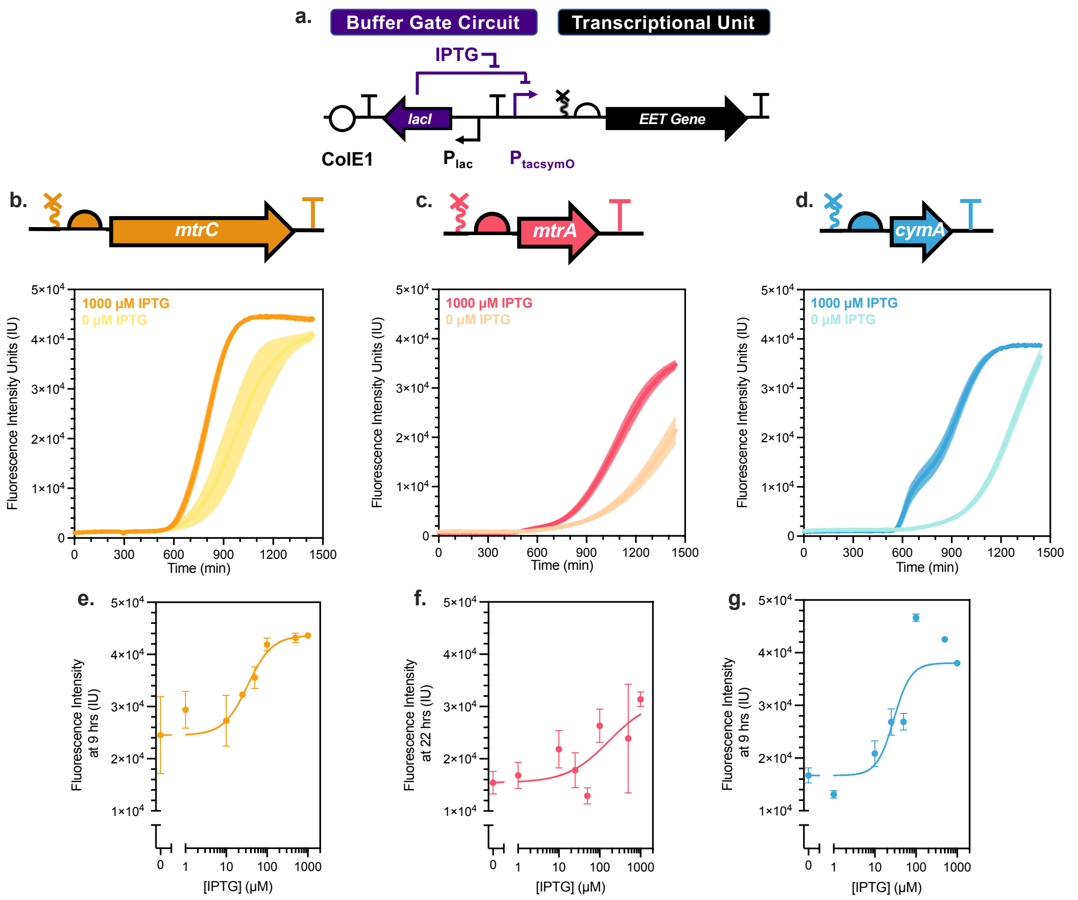


**Figure S11. a.** Diagram of generic Buffer gate circuit used to control gene of interest (*mtrC, mtrA,* or *cymA*) output in *S. oneidensis* JG596. CuAAC between 0.6 μM CalFluor 488 and 100 μM alkyne-PEG4-Acid in SBM. Raw aerobic kinetic curves for **b.** JG596 + pCD24r1 with *mtrC* inducible plasmid for both fully induced and uninduced kinetics inoculated at an $OD_{600}$ = 0.025 (5 x $10^7$ CFU/mL), **c.** *S. oneidensis* Δ*mtrA* + pCD25r0 with *mtrA* inducible plasmid for both fully induced and uninduced kinetics inoculated at an $OD_{600}$ = 0.025 (5 x $10^7$ CFU/mL), and **d.** *S. oneidensis* Δ*cymA* + pCD26r1 with *cymA* inducible plasmid for both fully induced and uninduced kinetics inoculated at an $OD_{600}$ = 0.025 (5 x $10^7$ CFU/mL). Hill function model for **e.** *S. oneidensis* JG596 + pCD24r1 **f.** *S. oneidensis* Δ*mtrA* + pCD25r0 and **g.** *S. oneidensis* Δ*cymA* + pCD26r1**.** Data show mean ± SD of n=3 independent experiment.

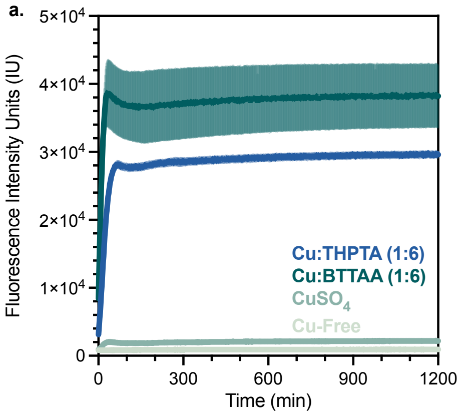


**Figure S12.** CuAAC between 0.6 µM CalFluor 488 and 100 µM alkyne-PEG$_4$-Acid in SBM. Aerobic reaction kinetics with sodium ascorbate and various ligands (300 µM).

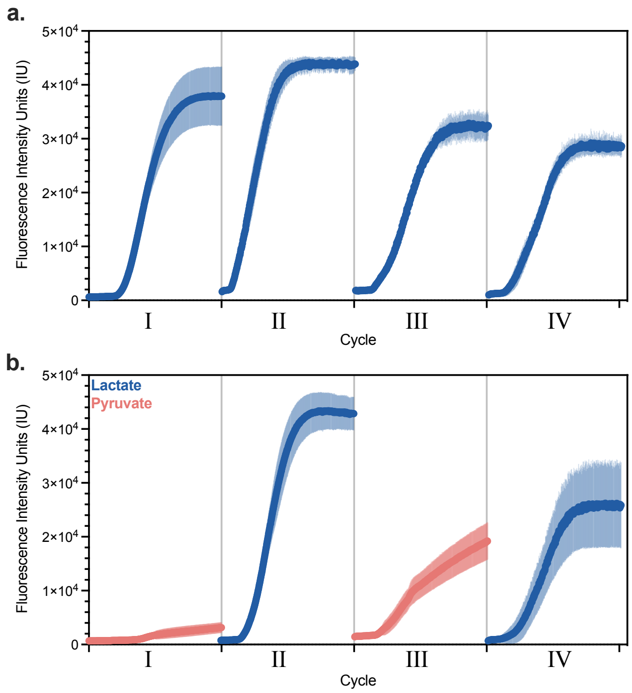


**Figure S13.** CuAAC cycling raw kinetic curves. **a.** Repeat CuAAC kinetics utilizing the same batch of *S. oneidensis* for each 11-hour cycle. **b.** Repeat CuAAC kinetics utilizing the same batch of *S. oneidensis* with different carbon sources for each 11-hour cycle. Data show mean ± SD of n=3 replicates.

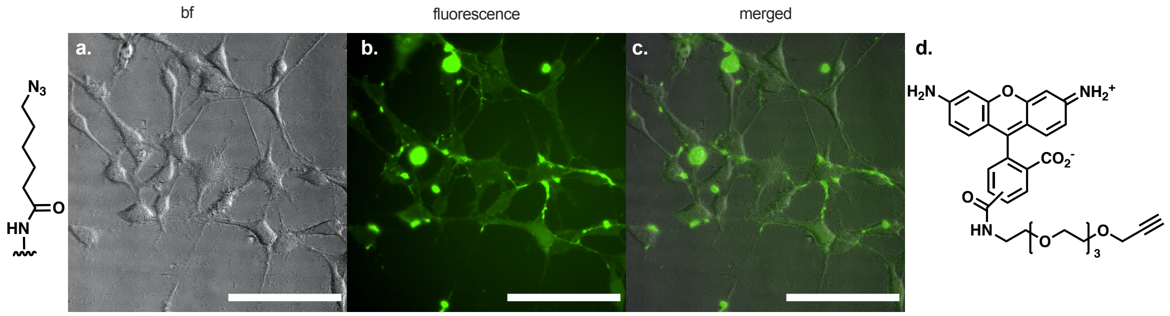


**Figure S13.** CuAAC performed in the presence eukaryotic cells **a.** Surface functionalization with 6-azidohexanoic acid (a terminal azide) through NHS ester displacement by free amines reacted for 3 h with carboxyrhodamine 110 in the presence of *S. oneidensis:* **c.** bright field images, **d.** fluorescent image, and **e.** merged image. Scale bars indicate 100 µm.
